# Rapid label-free identification of pathogenic bacteria species from a minute quantity exploiting three-dimensional quantitative phase imaging and artificial neural network

**DOI:** 10.1101/596486

**Authors:** Geon Kim, Daewoong Ahn, Minhee Kang, Jinho Park, DongHun Ryu, YoungJu Jo, Jinyeop Song, Jea Sung Ryu, Gunho Choi, Hyun Jung Chung, Kyuseok Kim, Doo Ryeon Chung, In Young Yoo, Hee Jae Huh, Hyun-seok Min, Nam Yong Lee, YongKeun Park

## Abstract

The healthcare industry is in dire need for rapid microbial identification techniques. Microbial infection is a major healthcare issue with significant prevalence and mortality, which can be treated effectively during the early stages using appropriate antibiotics. However, determining the appropriate antibiotics for the treatment of the early stages of infection remains a challenge, mainly due to the lack of rapid microbial identification techniques. Conventional culture-based identification and matrix-assisted laser desorption/ionization time-of-flight mass spectroscopy are the gold standard methods, but the sample amplification process is extremely time-consuming. Here, we propose an identification framework that can be used to measure minute quantities of microbes by incorporating artificial neural networks with three-dimensional quantitative phase imaging. We aimed to accurately identify the species of bacterial bloodstream infection pathogens based on a single colony-forming unit of the bacteria. The successful distinction between a total of 19 species, with the accuracy of 99.9% when ten bacteria were measured, suggests that our framework can serve as an effective advisory tool for clinicians during the initial antibiotic prescription.

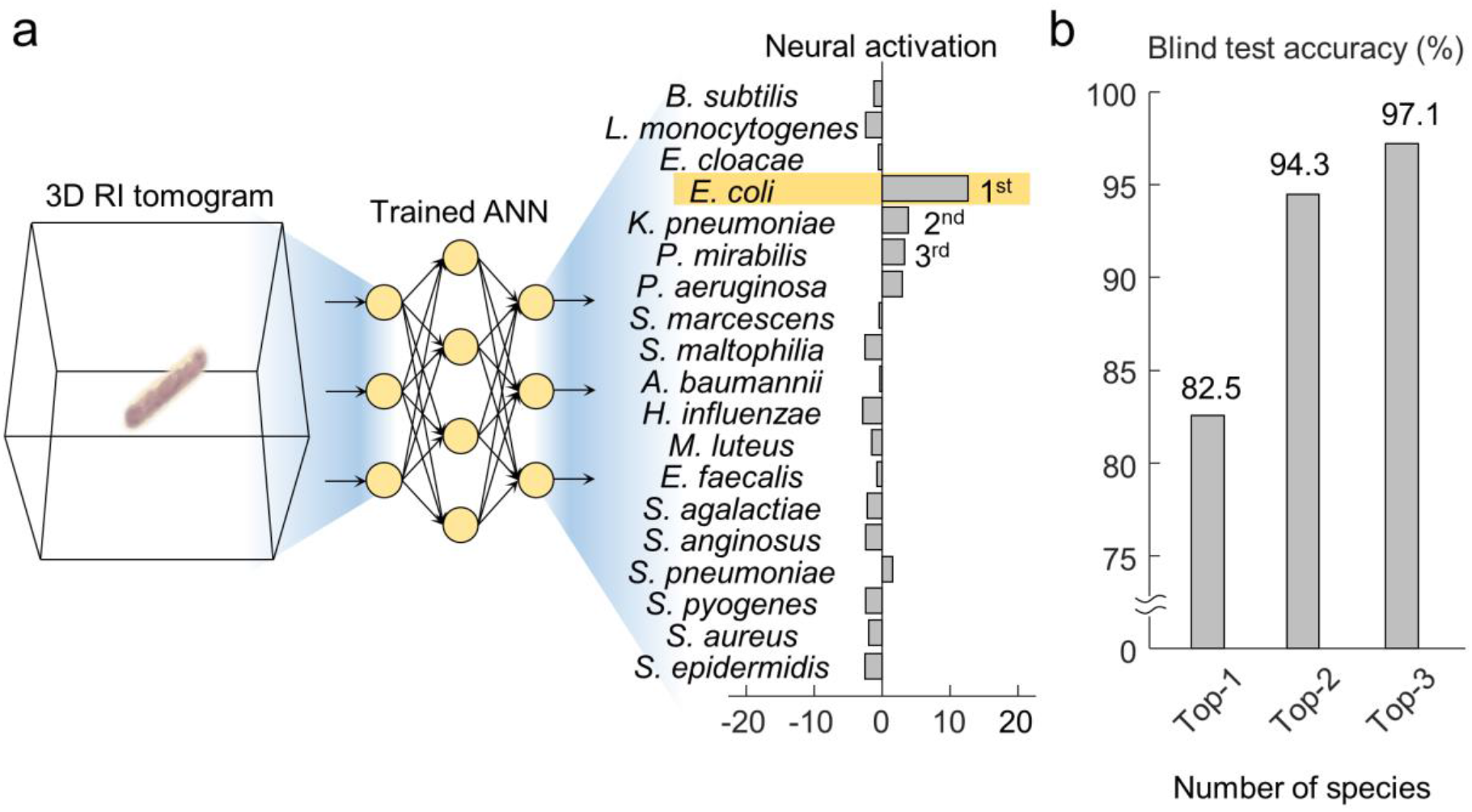

## 1. Introduction

Infection by microorganisms is one of the major healthcare issues worldwide, causing a significant number of casualties and a large amount of healthcare expense. Bacteria notably account for a large portion of life-threatening infections. During the year 2015, bacterial infections caused 4.4 million deaths, among a total of 8.8 million casualties by infections of any etiology (Hessling et al., 2017). In addition, the cost for treating bacterial infections accounts for 8.7% of the national health spedning in US (Torio and Moore, 2016).

The ideal treatment for an infection is the administration of appropriate antibiotics during the early stage. However, this is not easily implemented in the clinical settings, owing to the difficulty in rapid determination of the pathogen. Early prescriptions of antibiotics are commonly carried out empirically without the complete understanding of the etiology, and thus are often imperfect (García, 2009; Peterson et al., 2014) as antibiotics vary in the efficacy for different pathogens (Hutchings et al., 2019). A systematic review underlines that 46.5% of sepsis patients were given inappropriate empricial antibiotic treament and suffered 1.6-fold increased mortality risk (Paul et al., 2010). Accordingly, a rapid method for identifying the pathogen is required.

Conventional phenotypic approaches are time-consuming and often nonspecific, despite being relatively simple to perform (Bizzini and Greub, 2010). Culture tests, biochemical tests, and microscopic examination of gram-stained specimens are well-known conventional methods for microbial identification. They require hours or days of incubation for the metabolic activity or growth to take place. Molecular diagnostic methods are not scalable because of their process-specific sensitivity and high cost, even though they provide detailed information (Bizzini and Greub, 2010). Notably, 16S ribosomal RNA sequencing and real-time polymerase chain reaction offer genetic evidence regarding the identity of the pathogen. While these methods can precisely screen for a specific pathogen, the effectiveness of detection relies on the experimental setting, such as the choice of the primer or probe. Along with the relatively high cost, this technical intricacy limits the applicability of the molecular diagnostic methods.

In recent years, matrix-assisted laser desorption/ionization time-of-flight mass spectroscopy (MALDI-TOF MS) has become the gold standard for microbial identification (Bizzini and Greub, 2010; Seng et al., 2009), owing to its robust capability to investigate the molecular profile of the specimen. However, MALDI-TOF MS typically entails a turnaround time above 24 h, since sample amplification must precede to secure a detectable level of signal (Lin et al., 2018). A previous study indicated that a minimum of 10^5^ colony-forming units (CFUs) are required for MALDI-TOF MS-based detection of bacteria (Barreiro et al., 2017). In clinical settings, this quantity is obtained after hours or days of *in vitro* culture, while the mortality risk increases at an alarming rate (Moehring et al., 2013). Although some studies have sought alternative protocols to reduce the turnaround time of MALDI-TOF MS (Köck et al., 2017; Lin et al., 2018), the standard time-consuming protocol remains the most reliable approach.

To tackle this challenge of early microbial identification, we exploit quantitative phase imaging (QPI) and machine-learning-based image classification. Due to the noninvasive nature, QPI has facilitated quantitative investigations of live cells (Park et al., 2018b). Among various biomedical studies, bacteria has been investigated with QPI during growth (Ahn et al., 2020; Mir et al., 2011), while optically controlled in the presence of eukaryotic cells (Kemper et al., 2013), and upon the treatments of antibotics (Oh et al., 2020). In recent years, machine learning has been introduced to QPI (Jo et al., 2018; Rivenson et al., 2019b), enabling diverse applications including virtual staining (Rivenson et al., 2019a), virtual molecular imaging (Jo et al., 2020; Kandel et al., 2020), improvement of image quality (Kamilov et al., 2015; Ryu et al., 2019; Ryu et al., 2021), and a variety cell type classification (Chen et al., 2016; Rubin et al., 2019; Siu et al., 2020; Yoon et al., 2017). One noteworthy study realized efficient screening for anthrax spores using a handheld two-dimensional (2D) QPI microscope and artificial neural network (ANN) (Jo et al., 2017).

Here, we propose an image-based framework for the identification of bacterial species, which is accurate even for single or a few bacterial cells. We exploit the single-cell profiling ability of (3D) QPI in synergy with the image recognition power of ANN. Our ANN, that extracts the rich and complex information delivered by 3D QPI, facilitates accurate identification of bacterial species.The proposed framework was tested using our laboratory-obtained database of isolates that includes major bloodstream infection (BSI) pathogens. BSI, which is defined as the presence of microbial pathogens in the bloodstream, is a morbid disease with a mortality rate of approximately 25% and incidence rate around 200 per 100,000 people (Bearman and Wenzel, 2005). We demonstrate the species identification from a single or a few cells of BSI pathogens using our framework, evaluate the performance in multiple perspectives, and deliberate on how to circumvent the error.

## 2. Materials and methods

### 2.1. Preparation of bacteria

The bacterial samples were cultured *in vitro* from frozen glycerol stocks. The frozen stock of each species was stored at −80°C and thawed at room temperature (25°C) before use. After thawing, the stock was inoculated into a liquid medium and stabilized for over an hour in a shaking incubator at 35°C. The stabilized bacteria were seeded in agar plates containing a suitable medium. The agar plates were incubated at 35°C for 12-24 h until colony growth was visible. A liquid subculture seeded from the agar plate was incubated at 35°C for 8–16 h in a shaking incubator until the medium turned turbid. The subculture solution was diluted with a liquid medium to a concentration suitable for imaging, then sandwiched between cover glasses. Each species was inoculated in one of the following media: nutrient agar, brain heart infusion agar, tryptic soy agar, and chocolate agar. The glycerol stock or subculture was grown in nutrient broth, brain heart infusion broth, tryptic soy broth, or Giolitti-Cantoni broth.

### 2.2. Three-dimensional QPI

Each 3D refractive index (RI) tomogram was acquired using a commercialized 3D QPI system, also known as holotomography or optical diffraction tomography (ODT), (HT-2H, Tomocube Inc., Daejeon, Republic of Korea). The optical components are shown in Fig. 1(a). The system exploits Mach-Zehnder laser interferometry equipped with a digital micromirror device (DMD). As an optical analogous to X-ray computed tomography, ODT reconstructs the 3D RI tomogram of a transparent microscopic object from multiple 2D measurements of holographic images obtained with various illumination angles (Kim et al., 2016; Wolf, 1969). A continuous-wave laser with a wavelength of 532 nm serves as the light source. Two water-immersion objective lenses with 1.2 numerical aperture magnifies and de-magnifies the light.

**Fig. 1.**
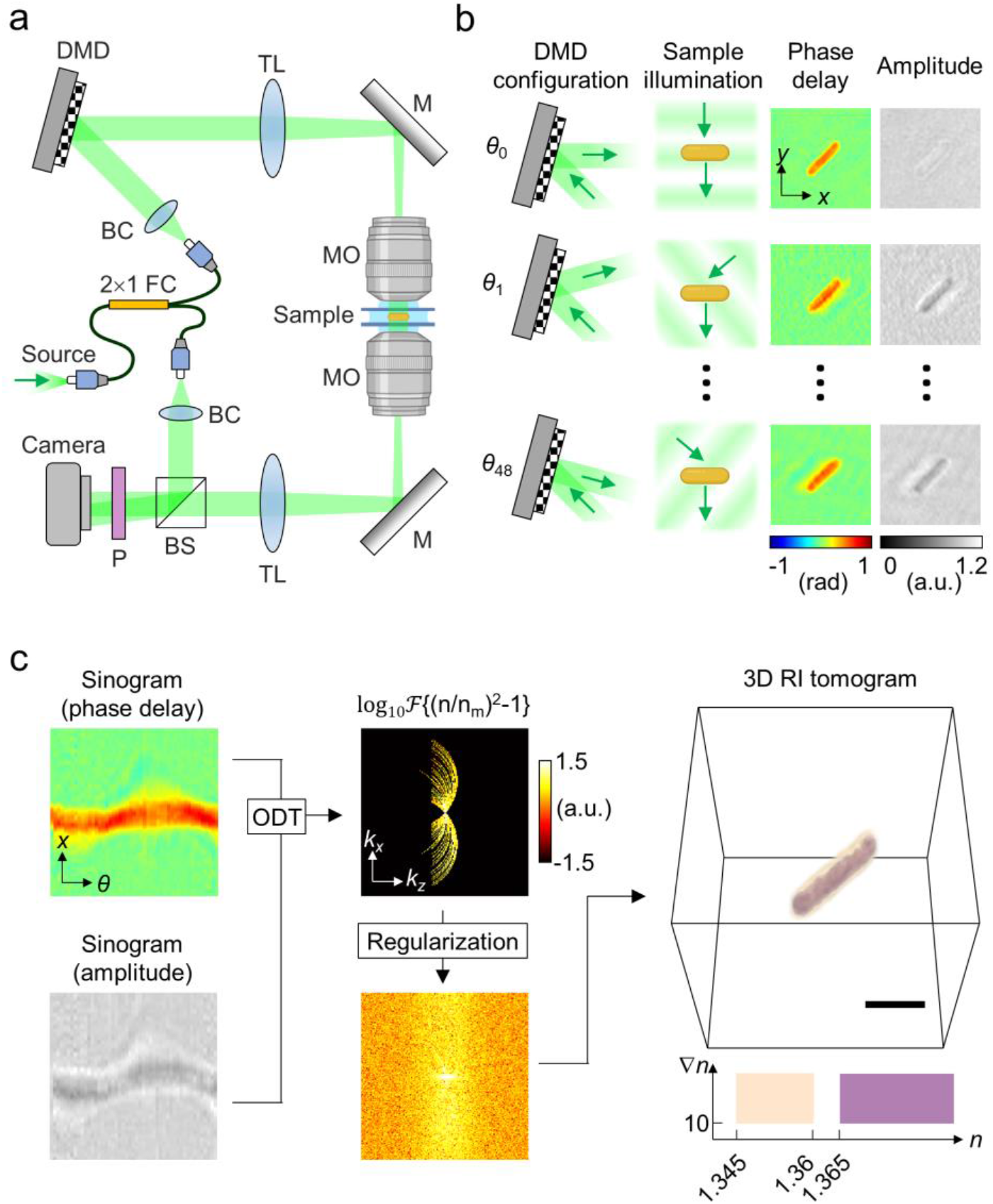
Three-dimensional quantitative phase imaging of bacteria. (A) The optical system is based on a simplified Mach-Zehnder interferometer. The incident angle of the light illuminated into the sample is controlled using the digital micromirror device, that is, utilizing the +1st order beam diffracted by the grating pattern on the DMD. BC: beam collimator, BS: beam splitter, CL: condenser lens, DMD: digital micromirror device, FC: fiber coupler, LP: linear polarizer, OL: objective lens, TL: tube lens. (B) Holograms including both the phase delay and the amplitude is measured while altering the illumination angle using the DMD. (C) The three-dimensional RI tomogram is acquired by integrating the sinogram into the scattering potential via optical diffraction tomography, followed by an iterative regularization.

The optical phase delay and the amplitude of light are retrieved from the 2D holographic image at each illumination angle (Fig. 1(b)), resulting in a sinogram of phase delay and amplitude. Here, the DMD alters the illumination angle by serving as a controllable binary grating (Lee et al., 2017; Shin et al., 2015). Our measurement scheme involves a total of 49 illumination angles. Once the sample is located in the microscope field-of-view, the 49 holographic measurements take approximately 0.4 sec. The 3D RI tomogram is reconstructed from the sinogram by inversely solving the Helmholtz equation followed by an iterative regularization (Lim et al., 2015) (Fig. 1(c)). The theoretical resolutions of the tomogram is 110 nm in the horizontal direction and 330 nm in the vertical direction, considering the spatial frequency range of the imaging system (Park et al., 2018a). Further descriptions regarding the computational details of ODT including the reconstruction algorithm can be found in previously published literature (Kim et al., 2013; Kim et al., 2016).

An individual 3D RI tomogram referred to the distribution of RI in a 12.8 μm × 12.8 μm × 12.8 μm volumetric space, sampled at a voxel resolution of 100 nm × 100 nm × 200 nm. Each 3D RI tomogram contained a single bacterium or several bacteria that were adherent together after fission; we term this qauntity a single CFU henceforth (1) in conformity to the definition of CFU (Hazan et al., 2012; Krieger, 2010) and (2) to connote the sample quantity required for our framework. A manual inspection of each 3D RI tomogram ensured that noisy measurements were ruled out before establishing the database.

### 2.3. Artificial neural network

The structure of ANN utilized in this framework mainly consists of 3D convolutional operations which can effectively explore the 3D structure of 3D RI tomograms. More specifically, the dense connections between the convolutional operations induce the ANN to revisit the feature maps of the shallower layers even at the deep layers. Fig. 2 illustrates the structure of our ANN in detail. The structure is inspired by the convolutional ANN design that outperformed most of the other designs in the benchmark tasks of 2D image analysis (Huang et al., 2017). The four dense blocks include 12, 24, 64, and 64 colvolutional operations, respectively, from shallow to deep. The number of feature channels after the initial convolution and the growth rate of the feature channels are 64 and 32, respectively.

**Fig. 2.**
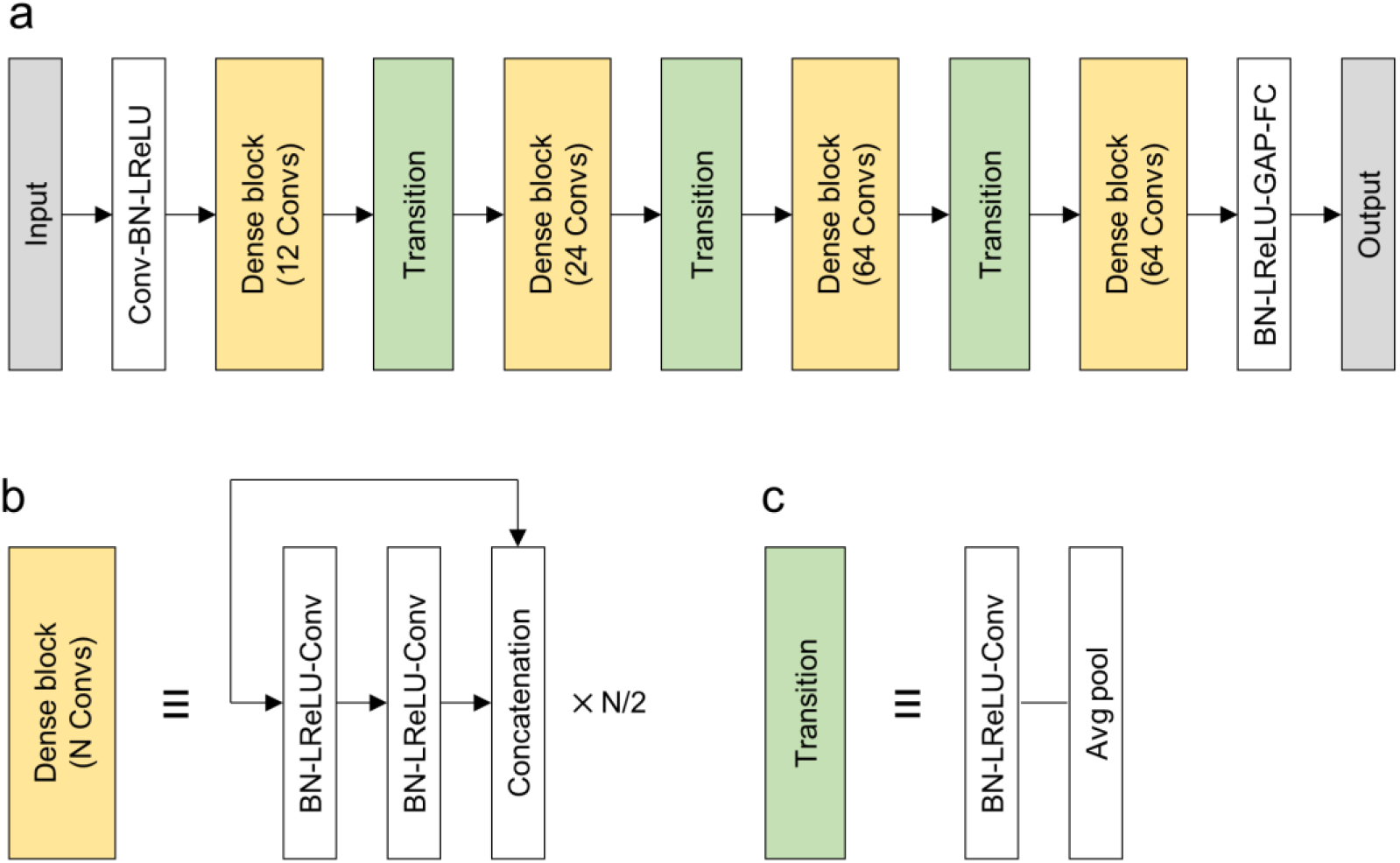
Structure of the artificial neural network. (A) The structure can be represented with four dense block transition units between adjacent dense blocks. Other elements include initial three-dimensional convolutional operation (Conv) of 3 × 3 × 3 kernels and a stride of 2 × 2 × 2, batch normalization, leaky rectified linear units, global average pooling, and fully connected operation. (B) The structure of the dense blocks allows features of shallower layers to be revisited in deeper layers. A dense block consists of a pair of Convs followed by concatenation of the feature map before the two Convs. In each pair of Convs, the first one has 1 × 1 × 1 kernels, and the second one has 2 × 2 × 2 kernels; meanwhile, the stride is 1 × 1 × 1 for both Convs. (c) The transition units shift the scale of the feature extracted by convolution. The Conv in each transition unit has 1 × 1 × 1 kernels and 1 × 1 × 1 stride.

The ANN was optimized by minimization of the cross-entropy loss between the ground truth and the prediction. For each species, 40 tomograms were randomly chosen as the blind test dataset and another 40 tomograms were randomly chosen as the validation dataset. The remaining tomograms composed the training dataset, which were directly reflected in the loss minimization process. The loss that occured in the training dataset was reduced using the stochastic gradient descent algorithm. The step size of the stochastic gradient descent algorithm was scheduled according to the cosine annealing method (Loshchilov and Hutter, 2016) at an initial step size of 0.001 and a period of 64 epochs. During training, data augmentation took place for each tomogram, once every epoch, to prevent overfitting of the trained model. The augmentation included random processes of horizontal crop, horizontal rotation, and Gaussian noise. During the blind test, each input tomogram was horizontally cropped around the center to provide an identical dimension. These processes resulted in an input tomogram of 9.6 μm × 9.6 μm × 12.8 μm to be fed into the ANN. A single training epoch through the entire training dataset took approximately 10 min, when using eight graphics processing units of GeForce GTX 1080ti and a central processing unit of Xeon E5-2600. The ANN was trained for 2,000 epochs while saving models that yielded high training accuracies or validation accuracies. The ANN and the optimization were implemented using PyTorch 1.0.0.

The algorithm for the blind test involved the predictions of multiple best-performing ANN models. Models with the highest accuracies for the training and validation datasets were chosen and integrated, to exploit a wider variety of features and prevent model-by-model variance. In search of the optimal strategy for chosing and integrating multiple models, four relevant parameters were explored. These parameters included the number of integrated models weighting between the accuracies for the training validation dataset, whether or not to normalize the neural activation, and the formula for integrating the predictions by the chosen models. Four options were considered as the formula for integrating the predictions: taking the average, taking the exponential average, voting, and taking the maximum projection of the neural activation. The combination of the parameters which yielded the highest validation accuracy established the algorithm for the blind test.

## 3. Results

The key function of our framework is to assess the species of the bacterial pathogen under a quantity of a single CFU level. 3D QPI and ANN classification can provide preliminary results during the early stages of infections, whereas the results of gold standard methods will be available dozens of hours later. Incorporation of our framework into the gold standard routine is practicable since our framework operates with a minute quantity of bacteria without destroying nor chemically modifying the bacteria.

### 3.1. Three-dimensional images of the bacteria

A database, which comprised 10,556 3D RI tomograms, was established with 19 different species of BSI pathogens. The 19 species accounted for around 90% of all BSI-related cases, as indicated by the annual data from a 1,000-bed tertiary care institute (Opota et al., 2015). The 3D RI tomograms of the 19 species showed that 3D QPI effectively conveys the microscopic structure of bacteria (Fig. 3). Some characteristic structures are clearly visible in the 3D RI tomograms, e.g., cellular chains of streptococci. The species and the corresponding numbers of tomograms are as follows: *Acinetobacter baumannii* (664), *Bacillus subtilis* (515), *Enterobacter cloacae* (541), *Enterococcus faecalis* (526), *Escherichia coli* (600), *Haemophilus influenzae* (511), *Klebsiella pneumoniae* (525), *Listeria monocytogenes* (632), *Micrococcus luteus* (247), *Proteus mirabilis* (517), *Pseudomonas aeruginosa* (596), *Serratia marcescens* (519), *Staphylococcus aureus* (558), *Staphylococcus epidermidis* (559), *Stenotrophomonas maltophilia* (549), *Streptococcus agalactiae* (537), *Streptococcus anginosus* (644), *Streptococcus pneumoniae* (566), and *Streptococcus pyogenes* (750).

**Fig. 3.**
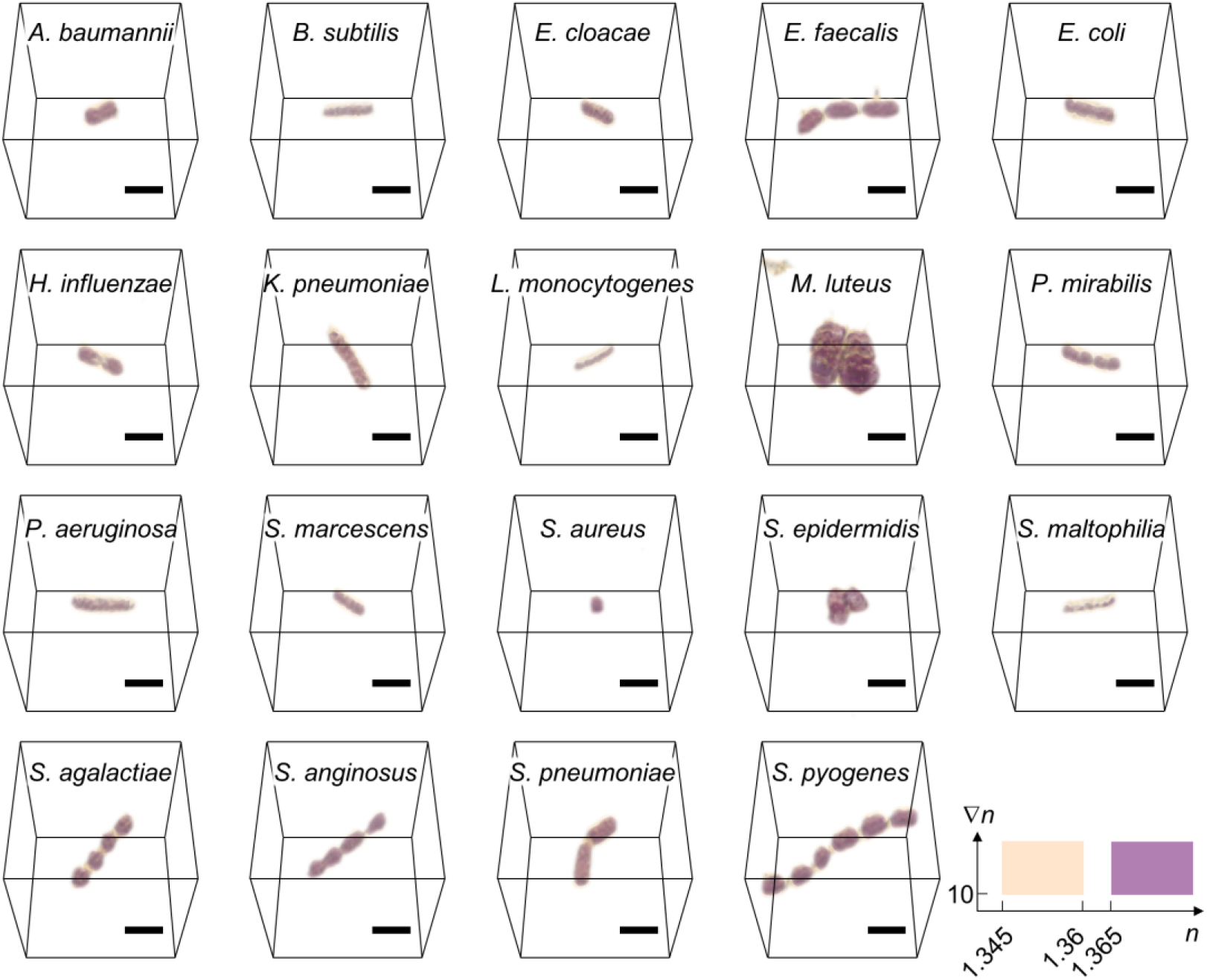
Three-dimensional refractive index tomograms of bacterial bloodstream infection pathogens. Representative tomograms addressed in out study are rendenred in three dimension. Each tomogram represents an individual species of bacterial pathogens. Scalebar = 2μm

### 3.2. Identification with single images

The optimized ANN model determined the species from an unseen 3D RI tomogram with an accuracy of 82.5% in the blind test. Our single-CFU accuracy was comparable to the rate of correct species identification using MALDI-TOF MS (Drancourt, 2010). During the training, the neural network was prompted to recognize structural features in the 3D RI tomograms of the training set and relate the features to the species. As a result, the ANN amplifies the neural activation of the species that share structural features with a blind test tomogram, whereas it attenuatese the neural activation of those that display distinct structural features (Fig. 4(a)). The species scoring the highest neural activation was chosen as the prediction result.

**Fig. 4.**
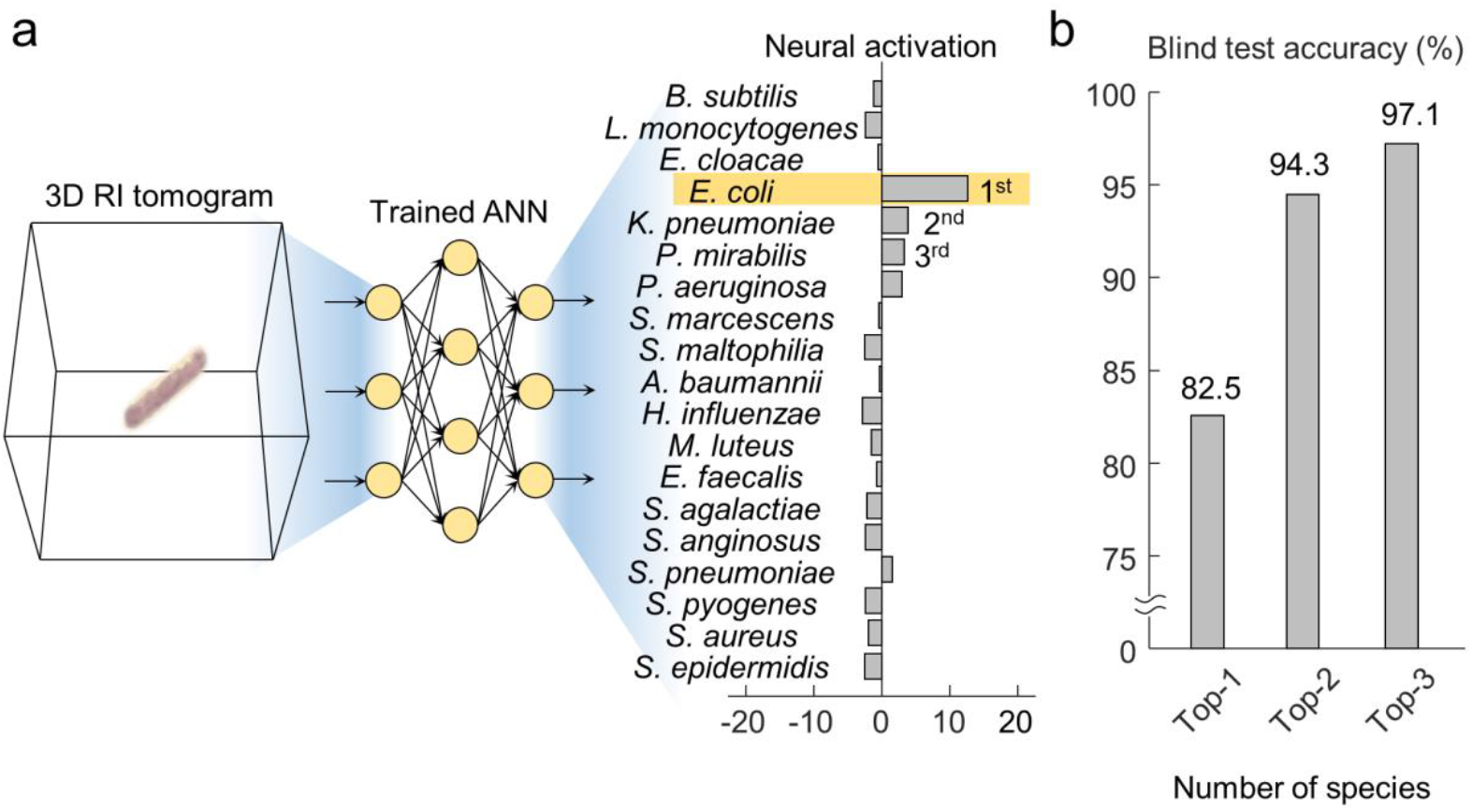
Identification of an individual three-dimensional (3D) refractive index (RI) tomogram using the artificial neural network (ANN). (A) The ANN model processes the given 3D RI tomogram and results in a neural activation profile, which represent the similarity in the space of features extracted by the ANN. (B) The resulting neural activation allows the accurate narrowing down of possible species and the selection of the single most likely species.

Examinations verified the significant roles of 3D QPI and ANN in achieving species identification at a single CFU level. 3D QPI presented the 3D structure of bacterial cells in detail, while the ANN extracted the underlying features from the complex domain of 3D RI tomograms. Our comparative study (Appendix A.1) showed that neither a 2D QPI measurement nor a multi-angular sinogram of 2D QPI measurements could reproduce the accuracy of 82.5%. In addition, our ANN outperformed the classification method based on machine learning and handcrafted features (Yoon et al., 2017) with more than twofold accuracy (Appendix A.2)

The risk of error could be reduced through a broader interpretation of the neural activaton. To be precise, narrowing down a few species that display high neural activation achieved lower rate of missing the correct species, compared to the single-species prediction. This approach secures additional sensitivity at the cost of specificity, which is a strategic trade-off. Approximately 94.3% of the blind test data included the correct species, two of which had the highest values of neural activation. The probability further increased to 97.1% when considering the top three values of neural activation (Fig. 4(b)). This considerable reduction of error was due to the robust feature-extracting ability of our ANN; the ANN recognizes the features related to the correct species, even in the cases of misidentified tomograms.

### 3.3. Error in species identification

To characterize the errors in each species, the blind test results for all 19 species were investigated using the confusion matrix (Fig. 5(a)). The most frequent errors included misidentification of *Acinetobacter baumannii* as *Streptococcus pneumoniae*, *Klebsiella pneumoniae* as *Streptococcus pneumoniae*, *Streptococcus agalactiae* as *Staphylococcus aureus*, and *Listeria monocytogenes* as *Bacillus subtilis*. Notably, thick bacilli and coccabacilli were prone to the misidentification as *Streptococcus pneumoniae*. This tendency is in agreement to the relatively elongated morphology of *Streptococcus pneumoniae* compared with other species of cocci (Hoyer et al., 2018; Pathak et al., 2018),

**Fig. 5.**
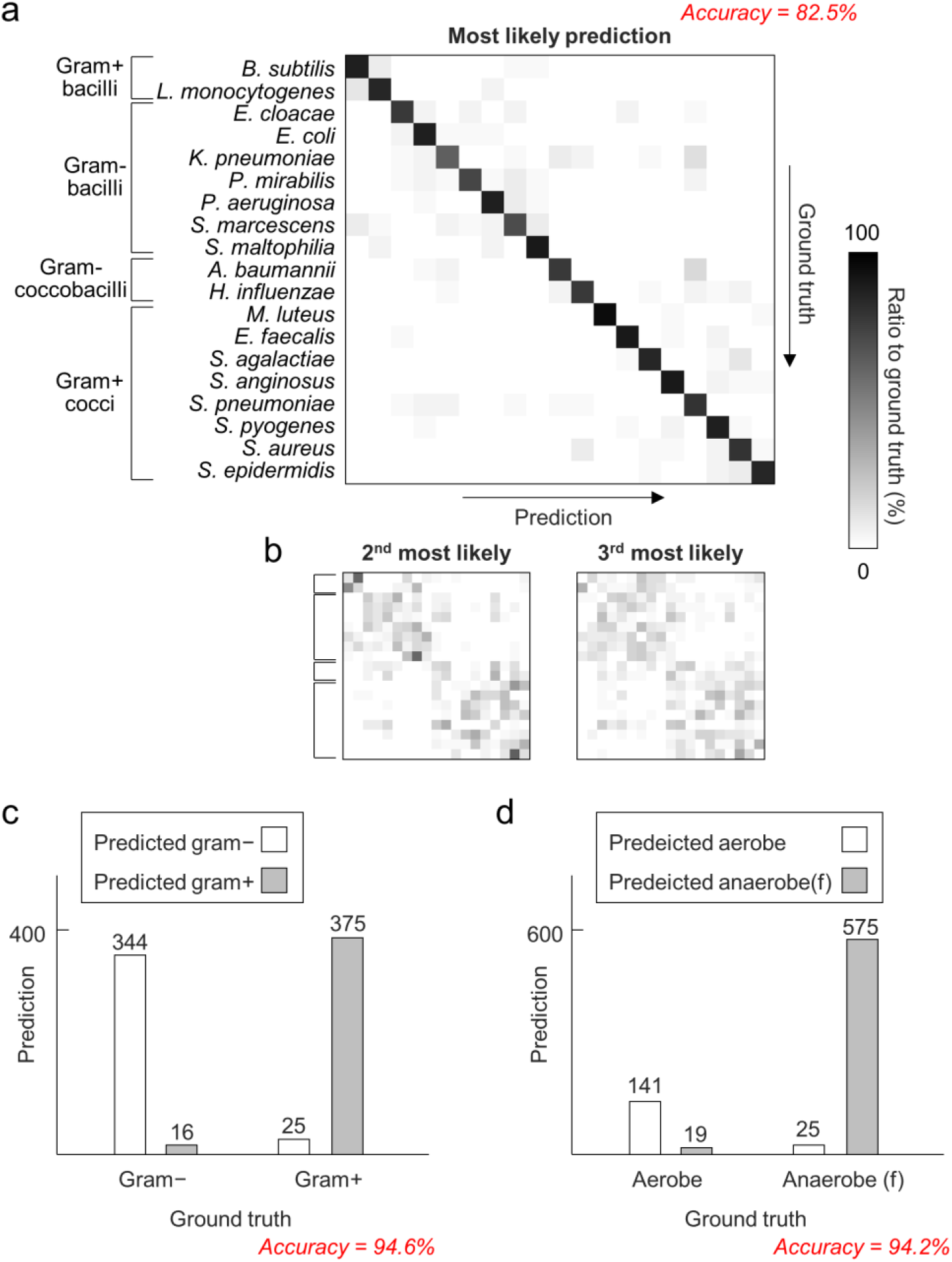
Distribution of error in species identification based on a single tomogram. (A) The confusion matrix visualizes the overall performance of the entire blind test dataset. The row and column indices correspond to the ground truth and the prediction, respectively. The indices of the 19 species are ordered to reflect the common bacterial categories. (B) The distribution of the second and the third most likely species further visualizes the interspecific similarity recognized by the trained artificial neural network (ANN). (C), (D) Individual tomograms are categorized under broader groups including gram-stainability and respiratory metabolism using a modified ANN for each task.

The distribution of the species with the second and third highest values of neural activation visualizes the similarity projected by the ANN (Fig. 5(b)). In this distribution, it is evident that the ANN reflects the morphological similarities between different species belonging to close categories. For instance, two clusters representing bacilli and other bacteria can be outlined, whereas the similarity between gram-positive bacilli further stands out compared with the similarity among all bacilli.

Classification tasks referring to the gram-stainability and respiratory metabolism were also carried out, resulting in accuracies of 94.6% and 94.2%, respectively (Fig. 5(c) and (d)). The higher accuracy of classification tasks compared with species identification indicates that the proposed framework can be conducted with higher certainty in broader categories.

### 3.4. Identification with multiple images

The accuracy of species identification significantly increased when multiple 3D RI tomograms were reflected in the prediction. The neural activation averaged over the inferences of multiple tomograms displayed a high probability to indicate the correct species, even for cases where most individual tomograms were misidentified (Fig. 6(a)). The error rate dropped more sharply than a simple reciprocal function of the number of tomograms; 94.9% and 98.4% accuracy was achieved, respectively using two and three 3D RI tomograms (Fig. 6(b)). This dramatic gain in accuracy was attributable to the robustness of the trained ANN in extracting species-related features, which was described in Section 3.2. In addition, a quantitative analysis underpinned that the correct predictions were made with higher contrasts in the neural activation compared to the mispredictions (Appendix A.3). This analysis explained how the averaging process selectively promotes correct predictions while flattening the erroneous signals in the neural activation.

**Fig. 6.**
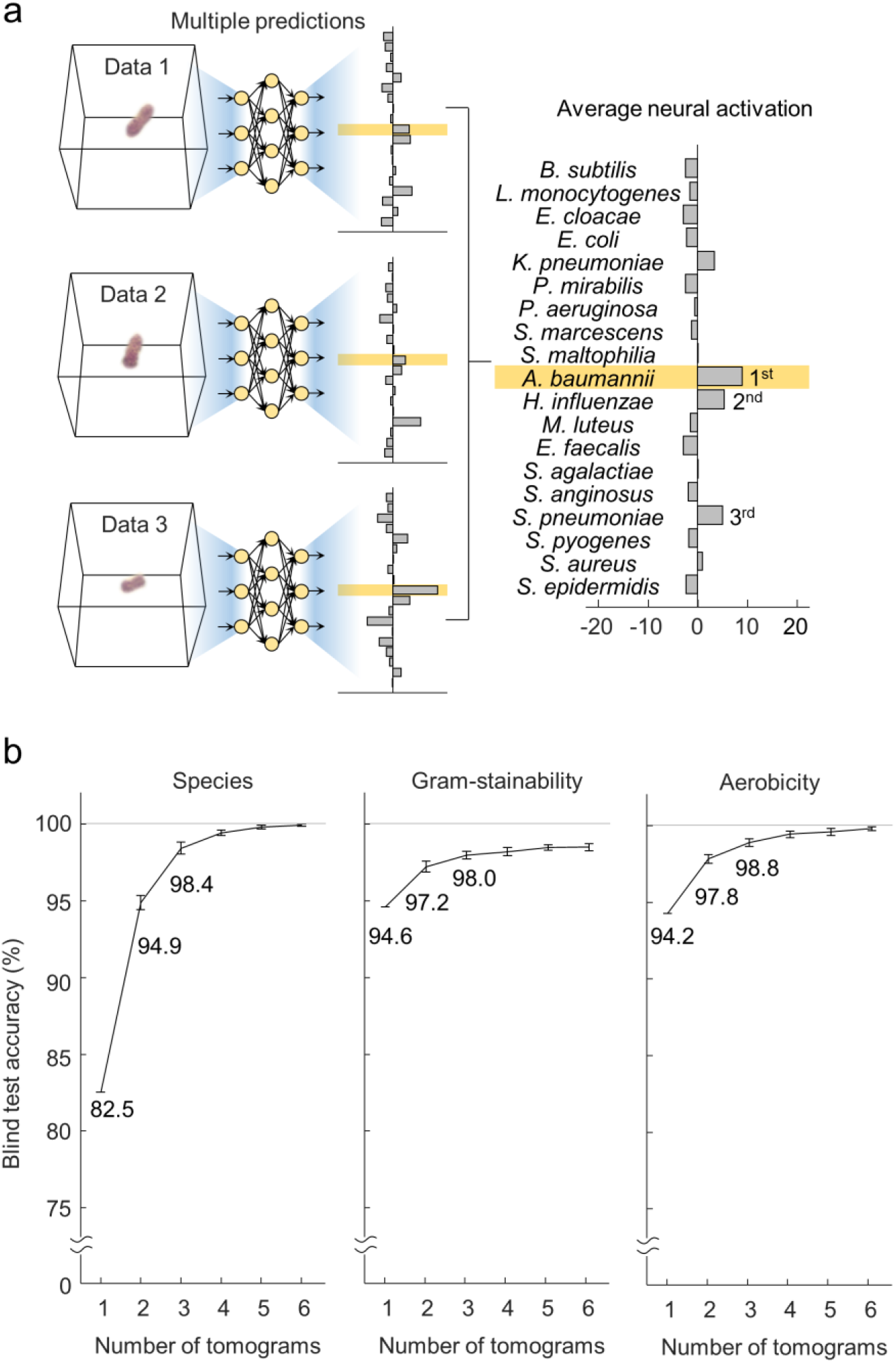
Increase o fidentification accuracy with multiple measurements. (A) A higher accuracy is obtained by taking the average of the neural activation resulting from multiple three-dimensional refractive index tomograms of an identical species. (B) The reduction of error is sharper than a simple reciprocal function owing to the feature-extracting ability of the artificial neural network.

## 4. Discussion

We propose a framework for species identification of bacterial pathogens at a single CFU level using 3D QPI and ANN. The exceptionally high accuracy under a limited sample quantity is attributable to the remarkable single-cell profiling ability of 3D QPI and the feature-extracting ability of ANN. Results show that the ample species-related features in a 3D RI tomogram are robustly extracted by the trained ANN, overcoming the quantity requirement of the previous methods.

We believe that the proposed framework will efficiently refine the initial antibiotics prescribed in the clinical settings. Our species identification accuracy based on a single CFU of bacteria is comparable to the performance of MALDI-TOF MS in identifying the species of blood-born pathogens (Drancourt, 2010). The risk of misidentification by our framework can be suppressed by taking multiple species into consideration. The risk of missing the correct pathogen dramatically dropped when two or three most likely species were selected. Our framework is also capable of being flexibly tuned for broader categories of bacteria such as gram-stainability or aerobicity. Even though these categories are not as specific as species, they can play a vital role in guiding antibiotic prescriptions. For instance, gram-positive pathogens can be effectively treated with vancomycin, whereas the gram-stainability can be determined by a destructive staining of sufficient sample. In addition, the accuracy of our framework can sharply increase through additional measurements of 3D RI tomograms. This signifies that our framework can be more accurate depending on the available sample quantity, compared to the baseline of the single-CFU performance. Furthermore, this framework can be incorporated along with the routine methods of microbial identification, including MALDI-TOF MS. The high performance at a minute sample quantity and the noninvasive property allow our framework to be added without exhausting the limited quantity of the sample.

Future studies on sample processing will propel our framework towards more immediate use. A condition suitable for imaging bacteria has to be met to perform our single-CFU level identification. For blood samples, this condition is achieved by performing lysis centrifugation after the initial blood culture (Kirn and Weinstein, 2013). However, application of our framework before completion of the blood culture is possible if a high-throughput procedure for enrichment of bacteria is introduced. A prominent and practical technique is the selective collection of particles utilizing advanced fluidic systems (Balyan et al., 2020; Kuntaegowdanahalli et al., 2009; Lee et al., 2019; Lei et al., 2012). Bacteria have also been prominent targets of collection using fluidic systems (D'Amico et al., 2017; Jung et al., 2020). The adoption of these strategies will facilitate the identification of the pathogen earlier than that suggested in our demonstration.

Moreover, validation using a larger diversity of pathogens will provide insights into the scope of application. We expect the proposed framework to be applicable to pathogens causing different classes of infections, such as urinary tract infections and lower respiratory infections, which are partially covered in this study. In addition, it is yet to be assessed whether the framework is capable of distinguishing strains resistant to antibiotics. The emergence of drug-resistant strains has compromised the established convention of antibiotic prescription, and the need to screen out resistant strains has also been highlighted (Chamieh et al., 2020; Hutchings et al., 2019; Shariati et al., 2020). Investigatig the performance in identifying bacterial strains and ensuring a higher accuracy to screening resistant strains will be crucial for improving the proposed framework.

## 5. Conclusion

To the best of our knowledge, our study demonstrates an unprecedented distinction of live and unmodified bacteria at a single CFU level among a wide range of species. With a single measurement of a bacterium or a CFU, we achieved the blind test accuracy of 82.5%, 94.6%, and 94.2% for species, gram-stainability, and aerobicity, respectively. With ten individual measurements, we achieved, the blind test accuracy of 99.9%, 98.9%, and 99.9% for species, gram-stainability, and aerobicity, respectively. Our accuracy based on a single measurement, which is comparable to the identification rate of MALDI-TOF MS, is facilitated by the precise 3D measurement of bacteria through 3D QPI and the statistical utilization of the measurement through ANN. We believe that the proposed framework will substantially augment the early countermeasures against bacterial infections; identifying the pathogen without the delay of sample amplification can provide a shortcut for administrating the appropriate antibiotics. We note that, in principle, the application of our framework can be brought forward to briefly after the sample collection if integrated with an advanced sample processing techniques.

## Appendix A

## A.1. Suitability of 3D QPI for species identification of bacteria

The benefit of 3D QPI in identification of bacteria at single CFU level was verified in a comparative experiment. Our proposed framework was compared with two other approaches utilizing the 2D equivalent of our 3D ANN structure. One approach is trained to identify the species from 2D amplitude and phase delay maps, while the other is trained to identify the species from the sinogram composed of 2D amplitude and phase delay maps in multiple illumination angles. The approaches based on 2D and sinogram data achieved 67.6% and 68.0% blind-test accuracy respectively after training and model incorporation identical to that of the 3D ANN (Fig. A.1). The significant difference in the accuracy suggests that 3D holographic microscope offers features to ANN in a more ostensive manner.

**Fig. A.1.**
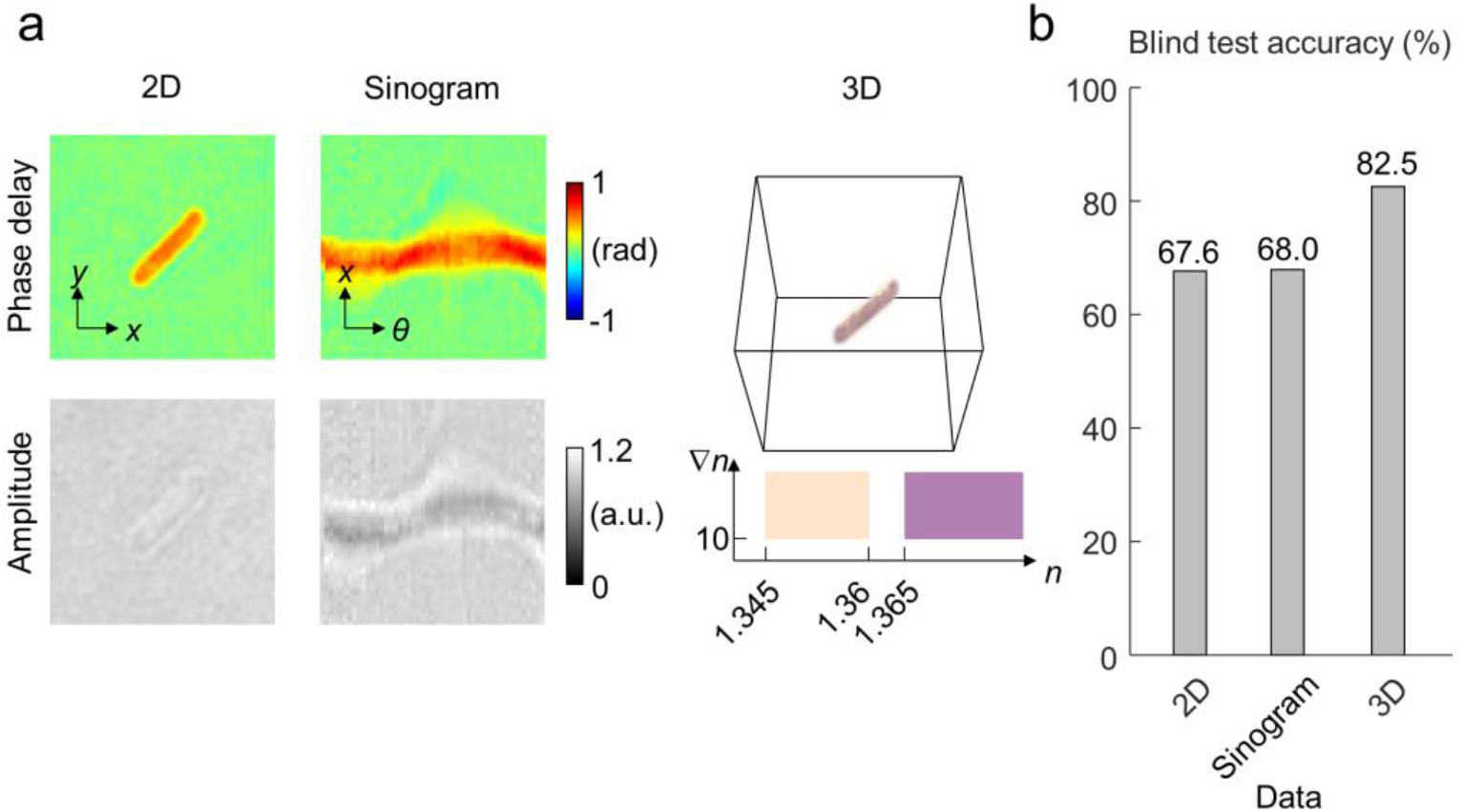
Comparison between three-dimensional (3D) and two-dimensional (2D) quantitative phase imaging (QPI) in image-based species identification. (A) Three types of QPI data are compared in the task of image-based species identification. A 2D QPI data consists of light phase delay and amplitude in a perpendicular illumination angle. A sinogram of 2D QPI data consists of light phase delay and amplitude in 49 illumination angles. The 3D QPI data refers to the 3D refractive index tomogram acquired from the 2D QPI sinogram. (B) The 3D QPI outperforms the 2D QPI and sinogram by a significant margin.

## A.2. Suitability of ANN for classification of 3D RI tomograms

The performance of ANN in recognizing the 3D RI tomograms of bacteria was compared to that of a conventional machine learning approach. For the comparison we apply the strategy of threshold-derived feature extraction and run *k*-nearest neighbors (*k*-NN) classifications based on the extracted features, that is, an approach that had effectively classified 3D RI tomograms of lymphocytes (Yoon et al., 2017) (Fig. A.2(a)). Scanning the values of *k* from 1 to 40, the lowest and the highest accuracies of the *k*-NN were 27.4% and 32.2% for *k* = 2 and *k* = 30 respectively (Fig. A.2(b)). As specified from the comparison, our implementation of ANN was more capable of recognizing species-related characteristics, compared to the machine learning with handcrafted features.

**Fig. A.2.**
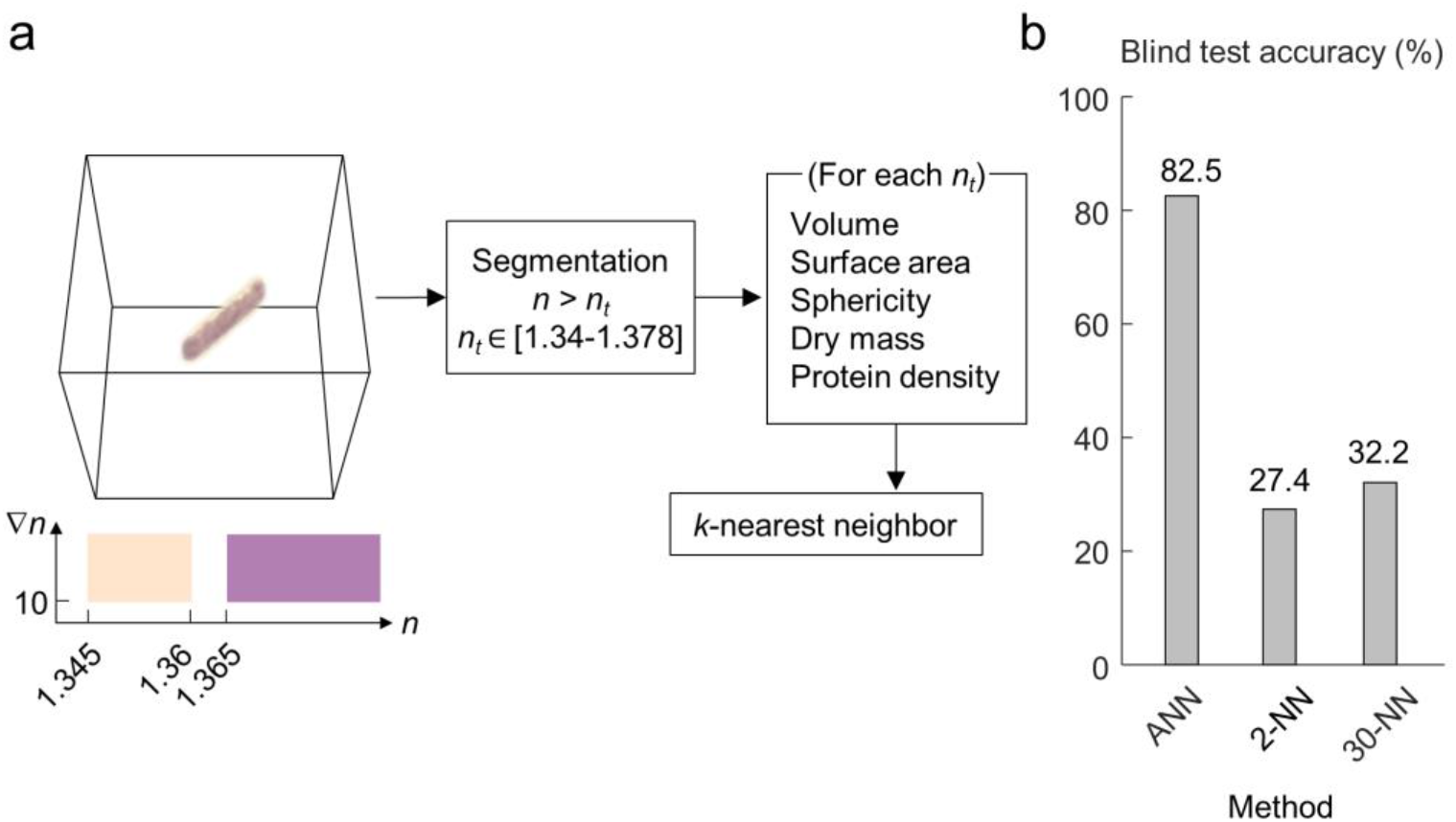
Comparison between artificial neural network (ANN) and conventional machine learning in image-based species identification. (A) An alternative identification approach based on handcrafted features of the three-dimensioanl refractive index tomogram and k-nearest neighbor (k-NN) algorithm. (B) The identification accuracy is compared between the proposed ANN approach and the k-NN analysis of handcrafted features. ANN outperforms k-NN by a huge margin, in both the best-performing case (k=30) and the worst-performing case (k=2).

## A.3. Contrast of neural activation

The dramatic rise of identification accuracy based on multiple 3D RI tomograms was accounted for by the feature-extracting ability of the ANN. A tendency appearing in the neural activation displays how the prediction significantly benefits from multiple tomograms. The contrast of neural activation, defined as the highest activation value divided by the sum of all other positive activation values (Fig. A.3(a)), was significantly higher in the correctly identified cases than the misidentified cases (Fig. A.3(b)). The difference displays how taking the average of neural activation from multiple 3D RI tomograms elevates the correct element of the neural activation.

**Fig. A.3.**
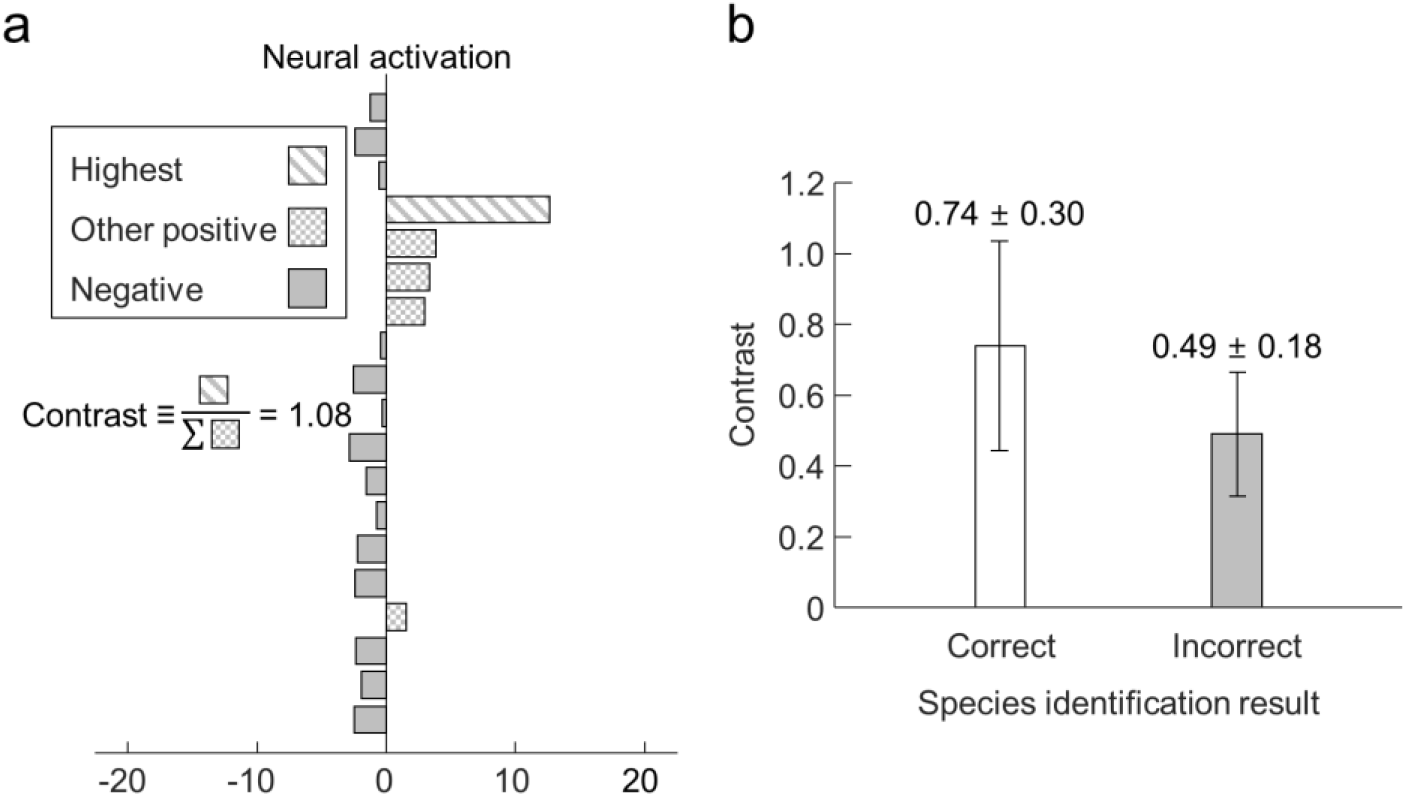
Contrast of neural activation for correctly and incorrectly identified tomograms. (A) The contrast of neural activation is defined as the highest neural activation divided by the sum of other positive neural activation, in order to represent the exclusiveness of the identification. (B) Statistical comparison of the identification contrast between the correctly identified test data and the misidentified test data.

## Acknowledgements

Y. Jo acknowledges support from KAIST Presidential Fellowship and Asan Foundation Biomedical Science Scholarship. The authors thank Ms. Hyeonjung Kim (The.Wave.Talk) for providing sample preparation protocols and agents. The clinical isolates were obtained from Asian Bacterial Bank of the Asia Pacific Foundation for Infectious Diseases.

## Author contributions

G.K., Y.J. and Y.P. conceived and designed the research. G.K., M.K., J.P., J.S., and J.S.R. conducted the experiments. G.K., D.A., G.C., and H.M. analyzed the data. G.K., Y.J., D.R., H.J.C., K.K., D.R.C., I.Y.Y, H.J.H, H.M., N.Y.L., and Y.P. prepared the manuscript. All authors read and discussed the results.

